# Enrichment of methylated cell-free placental DNA

**DOI:** 10.64898/2026.08.17.745276

**Authors:** Keaton W. Smith, Natalie Yuen, Shu Yi Shen, Sylvie Girard, Nicholas Cheng, Philip Awadalla, Tim Triche, Scott V. Bratman, Daniel D. De Carvalho, Elena Tuzhilina, Samantha L. Wilson, Michael M. Hoffman

## Abstract

**Introduction:** Preterm birth drives adverse perinatal maternal and infant health outcomes through heterogeneous symptoms, severity, and etiologies. Delivery prior to reaching 37 weeks of gestation may result from medically indicated intervention for pregnancy complications or spontaneously in the absence of prior symptoms. Placental tissue collected following preterm birth exhibits differential DNA methylation compared to full-term placentas and may indicate pregnancy health during gestation. Placental DNA currently has limited utility for assessing health of ongoing pregnancy, as sampling placental tissue during gestation increases the risk of infection and miscarriage. Risks associated with placental sampling during pregnancy limit the use of DNA methylation in clinical preterm birth prediction. Assessing preterm birth risk during gestation requires non-invasive methods for characterizing placental DNA methylation.

**Results:** We quantified genome-wide DNA methylation patterns of hypermethylated cell-free DNA in pregnant (*n* = 99) and non-pregnant (*n* = 93) plasma using cell-free methylated DNA immunoprecipitation sequencing (cfMeDIP-seq). In each sample, we assessed DNA methylation status in 300-bp genomic windows, examining both sequencing read counts and calculated absolute molar DNA amount. Known hypermethylated placental regions, including *RASSF1*, *STAT5A*, and *ERG* promoters showed significantly increased odds of detection in pregnant samples, suggesting enrichment of cell-free placental DNA. Of the 536,444 300-bp windows examined, 173,071 (32%) showed significant enrichment in pregnant plasma. Linear modeling identified 107,505 differentially methylated regions (DMRs) associated with pregnancies later diagnosed with intrauterine growth restriction (IUGR) (*n* = 22). Alu elements showed increased representation in these DMRs than expected, while other repetitive elements exhibited underrepresentation.

**Discussion:** These results demonstrate cfMeDIP-seq’s ability to enrich for cell-free placental DNA and characterize cell-free DNA methylation signatures of pregnancies complicated by IUGR. Enrichment of cell-free placental DNA enables non-invasive profiling of placental DNA methylation from maternal plasma. Detectable epigenetic signatures in maternal plasma may identify pregnancies at elevated risk for preterm birth before clinical symptoms appear. Our findings further highlight the potential of cell-free placental DNA for monitoring pregnancy health.

## Introduction

### Preterm birth

Preterm birth causes adverse outcome in short-term perinatal health and long-term maternal and infant health (Goldenberg et al. 2008). Defined as delivery prior to reaching 37 weeks gestational age, preterm birth occurs in approximately 11% of births globally (Goldenberg et al. 2008). It causes or contributes to an estimated 1 million infant deaths per year (Goldenberg et al. 2008; Bokslag et al. 2016; Armengaud et al. 2021). Roughly 30%–35% of preterm birth cases result from medical indication to manage pregnancy complications such as preeclampsia, intrauterine growth restriction (IUGR), or overt clinical infection through early delivery (Goldenberg et al. 2008). Most preterm birth consists of non-induced, idiopathic cases, with non-detectable infection or other unknown etiologies leading to preterm premature rupture of the membranes (PPROM) or spontaneous preterm labor with intact membranes (SPL) (Goldenberg et al. 2008; Goldenberg et al. 2000; Steegers et al. 2010; Lin et al. 2015; Suhag and Berghella 2013).

Symptoms and severity present highly heterogeneously within all subtypes of preterm birth (Moutquin 2003; Benton et al. 2018; Khan et al. 2023). Despite prior research on molecular, mechanistic, and epidemiological characteristics of preterm birth and its subtypes (Goldenberg et al. 2008; Knijnenburg et al. 2019), identifying at-risk pregnancies and accurately predicting the occurrence of preterm birth remains challenging.

### Placental dysfunction as a driver of preterm birth

A growing body of literature reports an association between poor placental health and preterm birth (Morgan 2016). During the first trimester of placental development, extravillous trophoblasts derived from progenitor cytotrophoblasts invade the maternal endometrium and remodel maternal spiral arteries (Cindrova-Davies and Sferruzzi-Perri 2022). This essential process establishes sufficient supply of maternal blood to the placenta, which facilitates the exchange of nutrients, gases, and waste between the mother and fetus (Cindrova-Davies and Sferruzzi-Perri 2022).

Insufficient spiral artery remodeling, placental injury, or placental inflammation can interfere with the supply of nutrients from the placenta to the fetus, causing placental insufficiency (Morgan 2016). Placental insufficiency can lead to both hypertensive disorders, such as preeclampsia, and poor growth conditions, such as IUGR (Cindrova-Davies and Sferruzzi-Perri 2022). Both of these outcomes contribute to medically indicated preterm birth (Cindrova-Davies and Sferruzzi-Perri 2022). Injury or inflammation of the placenta, such as chorioamnionitis, can also lead to idiopathic preterm birth via PPROM or SPL (Morgan 2016).

The placenta exhibits high adaptability and often undergoes functional and morphological changes to protect the fetus from potentially harmful environmental stimuli or conditions (Morgan 2016). For example, preeclampsia pregnancies can exhibit abnormally large or small placentas, while decreased placental size and thickness often occur in IUGR (Cindrova-Davies and Sferruzzi-Perri 2022; Dahlstrøm et al. 2008; Liu et al. 2021). Changes in placental size due to excessive or insufficient growth may indicate impaired placental function. Such impairment can lead to inadequate supply of nutrients and oxygen to the fetus. It remains unclear if placental phenotypic changes cause or compensate for preterm birth pathophysiology. Analysis of molecular changes may elucidate driving factors behind placental adaptation to environmental insults and offer insights into the etiology or preterm birth.

### DNA methylation and the placental methylome

Recent work uses of DNA methylation signatures to assess overall placental health and characterize molecular changes associated with preterm birth subtypes (Khan et al. 2023; Robinson and Price 2015; Wilson and Robinson 2018). DNA methylation refers to the addition of a methyl functional group to the carbon 5 of a cytosine base in a cytosine-guanine dinucleotide (CpG) site, forming 5-methylcytosine (Moore et al. 2013). As an epigenetic modification, DNA methylation can alter gene expression and regulation without changing the DNA sequence itself (Moore et al. 2013). DNA methylation plays an important role in regulating normal, cell-specific gene expression and can occur as a result of environmental stimuli (Moore et al. 2013).

The placental methylome, the genome-wide DNA methylation signature in the placenta, differs from somatic tissues and changes across gestation (Robinson and Price 2015). In early development, the preimplantation blastocyst adopts a characteristic global hypomethylated DNA methylation signature with erasure of most epigenetic modifications (Robinson and Price 2015). While epigenetic modification of DNA occurs extensively during fetal tissues differentiation, *de novo* DNA methylation remains infrequent across the placental genome (Robinson and Price 2015). CpG-dense genomic regions, such as gene promoters in CpG islands, become hypermethylated in trophoblasts and other placental cell types (Robinson and Price 2015). Some *de novo* methylation of placental DNA occurs normally throughout gestation to drive cell differentiation and adapt to stimuli. The placenta largely retains global hypomethylation with hypermethylated CpG-dense regions, which gives it a distinct DNA methylation signature compared to most non-placental tissues.

Transposable elements in the genome, such as long interspersed nuclear elements (LINEs), short interspersed nuclear elements (SINEs), and long terminal repeat (LTR) retrotransposons, regulate gene expression in the placenta through epigenetic mechanisms (Keighley et al. 2023). In somatic tissues, most transposable elements exhibit hypermethylation to silence their activity and prevent insertional mutagenesis (Keighley et al. 2023). Many transposable elements in the placenta exhibit global hypomethylation and act as alternative promoters and enhancers for trophoblasts (Keighley et al. 2023).

DNA methylation differs between preterm and full-term pregnancies in both maternal blood and placental tissue collected at delivery. (Brockway et al. 2023; You et al. 2021; Yuen et al. 2010; Ding 2012; Blair et al. 2013; Anton et al. 2014; Wilson et al. 2018; Banister et al. 2011; Ferreira et al. 2011; Koukoura et al. 2012; Wilhelm-Benartzi et al. 2012; Park et al. 2020; Keighley et al. 2023). Maternal blood from preterm pregnancies shows differential DNA methylation of genes involved in inflammation and immune pathways as well as EGFR, PRL, chemokine, interferon gamma, and Notch signaling (Knijnenburg et al. 2019). Dysregulation of placental transposable element DNA methylation appears in pregnancy complications, including preeclampsia and IUGR, suggesting a role in placental dysfunction (Keighley et al. 2023; Li et al. 2019; Wilhelm-Benartzi et al. 2012).

Many studies examining DNA methylation differences in placental tissue from preterm pregnancies either analyze preterm birth as a single homogenous condition. Failure to account for the heterogeneity of individual preterm birth subtypes may lead to potential masking of subtype-specific biological signals (Brockway et al. 2023; Yuen et al. 2010; Ding 2012; Blair et al. 2013; Anton et al. 2014; Wilson et al. 2018; Banister et al. 2011; Ferreira et al. 2011; Koukoura et al. 2012; Wilhelm-Benartzi et al. 2012). Several placental genes exhibit differential DNA methylation in idiopathic preterm birth relative to full-term pregnancies (Wang et al. 2019). Conflicting evidence suggests idiopathic preterm placentas often possess an overall similar DNA methylation profile to those from full-term births (Wang et al. 2019; Brockway et al. 2023). Placentas from preterm pregnancies show widespread changes, including differential DNA methylation at the promoters of genes involved in inflammation, cell adhesion, trophoblast differentiation, and cellular metabolism (Yuen et al. 2010; Ding 2012; Blair et al. 2013; Anton et al. 2014; Wilson et al. 2018). Similarly, genes belonging to angiogenic and growth factor pathways show differential DNA methylation in growth-restricted pregnancies compared to pregnancies with appropriate fetal growth (Banister et al. 2011; Ferreira et al. 2011; Koukoura et al. 2012; Wilhelm-Benartzi et al. 2012).

Heterogeneity and molecular differences within and between idiopathic and medically indicated preterm birth suggest the presence of distinct subtypes of disease. Many studies investigating DNA methylation profiles of placentas from preterm pregnancies do not account for the high degree of preterm birth heterogeneity, leading to conflicting reports of DNA methylation characteristics of preterm birth in the literature (Khan et al. 2023).

Placental DNA methylation studies usually examine placental tissue collected at the time of delivery, preventing assessment of pregnancy health during gestation. Investigating placental DNA methylation signatures prior to delivery could provide a novel way to monitor the risk of preterm birth before the onset of symptoms. Sampling placental tissue prior to delivery (known as chorionic villus sampling) remains non-routine in most pregnancies due to increased likelihood of infection, miscarriage, and preterm birth itself (Salomon et al. 2019). These risks currently serve as a major hindrance to the use of placental DNA methylation signatures in assessing pregnancy health.

### Cell-free placental DNA methylation as a non-invasive indicator of placental health

Cell-free DNA shed from the placenta into maternal circulation offers a non-invasive means of examining placental genetic and epigenetic characteristics such as genetic sequence and DNA methylation (Yuen et al. 2024). Cell-free placental DNA, sometimes called “cell-free fetal DNA”, refers to fragmented cell-free DNA present in the maternal circulation throughout pregnancy originating from normal trophoblast turnover via apoptosis or necrosis (Yuen et al. 2024). While most circulating cell-free DNA derives from maternal hematopoietic tissues, cell-free placental DNA accounts for 20%–30% of the total cell-free DNA by late gestation (Fan et al. 2008; Sun et al. 2015). Non-invasive prenatal testing uses cell-free placental DNA to detect fetal aneuploidies and other genetic conditions (Shear et al. 2023).

Cell-free placental DNA reflects the DNA methylation patterns of placental tissue and exhibits average global hypomethylation with hypermethylated CpG-dense regions (Lun et al. 2013; Jensen et al. 2015; Chu et al. 2021). Recent studies explore the utility of cell-free DNA methylation profiling to identify epigenetic markers of disease and determine DNA fragment tissue-of-origin (Sun et al. 2015; Li et al. 2023). Some epigenetic characteristics of placental DNA serve as markers for placental origin of cell-free DNA (Yuen et al. 2024). Distinct nucleosome positioning and DNA methylation characteristics such as hypomethylation of *PDE9A* and hypermethylation of the *RASSF1*, *STAT5A*, *ERG*, *HLCS*, and *UMODL1* gene promoters suggest placental origin (Yuen et al. 2024). Hypermethylated *RASSF1* promoter quantification allows clinical estimation of the overall percentage of cell-free DNA deriving from placental tissue, also called the “fetal fraction” of cell-free DNA (Chan et al. 2006; ’t Oever et al. 2025). Selectively analyzing placental-sourced cellfree DNA fragments by enriching for cell-free placental DNA from maternal blood offers a unique glimpse into the placental methylome during gestation (Del Vecchio et al. 2021). Characterization of genome-wide cell-free placental DNA methylation may allow non-invasive assessment of placental and overall pregnancy health (Del Vecchio et al. 2021).

In cancer, circulating tumour DNA allows detection and classification of solid tumours based on DNA methylation signatures (Diaz and Bardelli 2014). Similar to cell-free placental DNA, circulating tumour DNA features global low levels of DNA methylation with hypermethylated CpG dense regions (Novakovic and Saffery 2013). Cell-free methylated DNA immunoprecipitation sequencing (cfMeDIP-seq) uses 5-methylcytosine monoclonal antibodies to enrich for low concentrations of hypermethylated circulating tumour DNA (Shen et al. 2018; Shen et al. 2019; Wilson et al. 2022). cfMeDIP-seq has shown robust performance in detecting solid tumours and can accurately classify a range of early-stage cancers based on their cell-free DNA methylation signatures (Shen et al. 2018; Lu et al. 2022; Zuccato et al. 2023; Burgener et al. 2021; Lasseter et al. 2020; Grisolia et al. 2024). cfMeDIP-seq specifically enriches for hypermethylated CpG-dense circulating tumour DNA fragments, enabling sensitive detection of low-abundance circulating tumour DNA in blood. The shared characteristics of tumour and the placental DNA, global hypomethylation and CpG dense region hypermethylation, suggest the use of cfMeDIP-seq for non-invasive assessment of placental health during pregnancy. Cell-free placental DNA levels in blood typically exceed those of circulating tumour DNA by approximately 10-to 100-fold (Bryzgunova et al. 2021; Cheng et al. 2021; Ashoor et al. 2013; Bischoff et al. 2005). This suggests that cfMeDIP-seq could readily detect cell-free placental DNA in maternal blood. Just as distinct circulating tumour DNA hypermethylation patterns distinguish solid tumour subtypes, cell-free placental DNA hypermethylation patterns in maternal circulation could provide an early identifier of preterm birth risk. Enrichment of placental-derived fragments may increase detection of placenta-specific DNA methylation signatures without eliminating potentially informative DNA methylation signal from maternal tissues. Leveraging the similarities between cell-free placental DNA and circulating tumour DNA methylomes, we hypothesized that cfMeDIP-seq enriches for cell-free placental DNA in maternal circulation and may detect DNA methylation signatures indicative of preterm birth and its subtypes.

Here we show that cfMeDIP-seq enriched for second trimester cell-free placental DNA in maternal plasma. Furthermore, we show that cfMeDIP-seq can characterize cell-free DNA methylation signatures of preterm birth subtypes, particularly IUGR, prior to symptom onset. These findings highlight the potential use of cfMeDIP-seq for non-invasive assessment of placental health and early identification of pregnancies at risk for preterm birth.

## Methods

### Sample collection

We obtained second trimester maternal plasma samples (*n* = 172) from the UK Baby Biobank (*n* = 83; University College London, Leon et al. 2016), Building Blocks of Pregnancy Biobank (*n* = 76; Indiana University), and the Global Alliance to Prevent Prematurity and Stillbirth (*n* = 13; Seattle Children’s Hospital), selecting samples with approximately equal representation of maternal age, ethnicity, and fetal sex when possible. This included samples from pregnancies resulting in medically indicated preterm birth subtypes such as IUGR, preeclampsia occurring after 34 weeks gestation notated as late-onset preeclampsia (LOPE), and preeclampsia. Preeclampsia either occurring before 34 weeks gestation or LOPE co-occurring with IUGR both received annotations as placental preeclampsia (pPE) due to their strong association with placental insufficiency. We also included idiopathic preterm birth subtypes such as PPROM and SPL as well as samples from healthy full-term control pregnancies. We obtained non-pregnant female plasma samples (*n* = 97) from the Ontario Health Study biobank (Kirsh et al. 2023).

### Cell-free methylated DNA immunoprecipitation sequencing

The Ontario Institute of Cancer Research performed cfMeDIP-seq on pregnant maternal plasma samples (*n* = 172) as previously described (Shen et al. 2019) with the addition of spike-in controls for methylated cell-free DNA quantification (Wilson et al. 2022). Cell-free DNA extraction used a QIAamp circulating nucleic acid kit (QIAgen, Hilden, Germany, cat. #55114). All samples contained at least 0.5 ng of extracted DNA prior to library preparation. DNA fragments received barcodes and adapters for sequencing. Pregnancy samples received 0.01 ng of synthetic spike-in control fragments, allowing for calculation of absolute molar amount of methylated DNA as per Wilson et al. 2022. Highly methylated DNA fragments isolated using 5-methylcytosine monoclonal antibodies (Diagenode, cat.#C15200081) underwent sequencing on an Illumina NovaSeq 6000 to obtain 60 million paired-end 100 bp reads per sample. A separate cohort of non-pregnant plasma samples (*n* = 97) from the Ontario Health Study biobank underwent cfMeDIP-seq following an identical protocol but without the addition of spike-in controls.

### Data preprocessing

We calculated total molar amount of methylated cell-free DNA for each fragment in the pregnant samples using the spike-in controls described by Wilson et al. 2022. We aligned fragments to GRCh38/hg38 (Schneider et al. 2017) using Bowtie2 (version 2.3.5.1; Langmead and Salzberg 2012) and divided reads into 300 bp windows (*ρ* = 543, 134) using bedtools (version 2.29.2; Quinlan and Hall 2010), as described by Shen et al. 2018 and Wilson et al. 2022. A generalized linear model based on read count, G+C content, CpG fraction, and fragment length approximated the absolute molar amount of DNA in picomoles from spike-in fragment recovery.

### Data filtering

To filter for high quality samples, we removed samples with fewer than 5 million unique cfMeDIP-seq reads or spike-in methylation specificity <95% (pregnant *n* = 73, non-pregnant *n* = 4). We excluded genomic windows overlapping regions in the ENCODE blacklist (*ρ* = 200; version 2, Amemiya et al. 2019) or with Umap multi-read-mappability ≤0.5 (*ρ* = 6, 490; Karimzadeh et al. 2018) as per Wilson et al. 2022. Sample and window filtering steps left 93 non-pregnant and 99 pregnant (control *n* = 22, IUGR *n* = 22, LOPE *n* = 23, PPROM *n* = 13, pPE *n* = 14, SPL *n* = 5) samples each with 536,444 genomic windows for analysis.

### Assessing cfMeDIP-seq’s ability to enrich for placental DNA

We used sample read counts to assess whether cfMeDIP-seq enriched for cell-free placental DNA. For each genomic region, non-zero read counts indicated the presence of detectable hypermethylated DNA fragments, while zero-read counts reflected fragments either below a detectable concentration or simply absent from a sample. As with many DNA enrichment protocols, genomic windows with zero read counts may represent DNA fragments present below a detectable concentration or truly absent from a sample, with no reliable way to distinguish one from the other. We considered genomic windows significantly more likely detected in pregnant compared to non-pregnant samples as enriched by cfMeDIP-seq in the pregnant dataset. We used negative binomial distributed zero inflation hurdle models (Cragg 1971; Mullahy 1986) to model the relationship between read count and pregnancy status (pregnant or non-pregnant). The hurdle models adjusted for body mass index (BMI), which can affect cell-free DNA concentration in plasma (Yuen et al. 2024). We fit these models for genomic windows surrounding regions known for hypermethylation in placenta and hypomethylation in somatic tissue: *RASSF1* gene promoter (chr 3: 50,340,550 – 50,342,500), *STAT5A* gene promoter (chr 17: 42,297,438 – 42,299,714), and *ERG* gene promoter (chr 21: 38,660,634 – 38,661,788) (Zhang et al. 2021; Chan et al. 2006; Rahat et al. 2016; Chen et al. 2015). These regions remain consistently hypermethylated in placental tissue and not methylated in other somatic tissues, and so we expected their detection by cfMeDIP-seq assuming successful cell-free placental DNA enrichment (Zhang et al. 2021; Chan et al. 2006; Rahat et al. 2016; Chen et al. 2015). We included a negative control region (chr 2: 240,085,545 – 240,101,752) known to exhibit stable levels of methylation across different tissue types (Edgar et al. 2014).

A Wilcoxon test assessed differences in the average number of reads for these regions between the pregnant and non-pregnant samples. In the pregnant samples only, we calculated Pearson correlation coefficients between gestational age and the total number of reads for each region of interest. For all tests, we considered a p-value <0.05 statistically significant.

We fit zero inflation hurdle models for each of the 300 bp genomic windows (*ρ* = 536, 444) available in the dataset. Calculating the probability of non-zero counts for a given window requires the window to have both zero and non-zero counts. Due to these models requiring a combination of zero and non-zero counts, we ignored genomic windows with zero read counts in all samples or non-zero read counts in all samples (*ρ* = 54, 028). We especially focused on the regions (*ρ* = ±11) surrounding *RASSF1*, *STAT5A*, and *ERG* promoters, with nearby genomic regions acting as negative controls where the literature reports no hypermethylation in placenta and we expected to see low or no enrichment. A false discovery rate (FDR)-adjusted p-value < 0.05 indicated statistical significance. Regions with a statistically significant increase in the odds of non-zero count in pregnant samples indicate higher likelihood of detectability in the circulation of pregnant individuals. We visualized the regions surrounding the transcription start sites (TSSs) of *RASSF1*, *STAT5A*, and *ERG* using genome browser plots produced in ggplot2 (version 3.5.1; Wickham 2016).

### Identifying differentially methylated regions in preterm birth subtypes

Using the limma package (version 3.58.1; Ritchie et al. 2015) for R (version 4.4.2; R Core Team 2024), a linear model identified differentially methylated regions (DMRs) between samples from each preterm birth subtype and healthy control pregnancies (Equation 1). We modeled picomolar amount of DNA (*Y*_i_) in a given genomic window as a function of gestational age (*x*_gestational age_), fetal sex (*x*_fetal sex_), maternal age (*x*_maternal age_) and BMI (*x*_BMI_), and the total number of cfMeDIP-seq reads (*x*_total reads_). The model optimizes regression coefficients (*β*) to adjust for each covariate and minimize residuals (*ε*_i_), as shown in Equation 1.

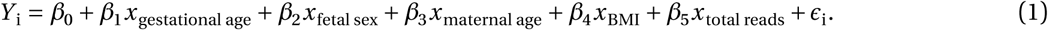

An FDR corrected p-value <0.05 indicated statistical significance.

### Assessing representation of transposable elements in differentially methylated regions

We annotated DMRs mapping to repetitive elements according to RepeatMasker and assessed over or underrepre-sentation of transposable elements. We randomly shuffled genomic windows from the full cfMeDIP-seq dataset and repeatedly (*X* = 1, 000 permutations) sampled sets of equal size to the DMR set (*ρ* = 107, 505). We reannotated each resampled set and estimated the expected distributions of LINEs, Alu elements, non-Alu SINEs, LTRs, and total repetitive elements. Given these distributions, we calculated the probability of the observed number of each element in the DMRs using the pnorm() function in R (version 4.4.2, R Core Team 2024).

## Results

### Cell-free methylated DNA immunoprecipitation sequencing enriches for hypermethylated cell-free placental DNA

The genomic windows surrounding the *RASSF1* (*ρ* = 4 windows), *STAT5A* (*ρ* = 5 windows), and *ERG* (*ρ* = 3 windows) promoters all showed a significant increase in the odds of non-zero read count in pregnant compared to non-pregnant cfMeDIP-seq samples (Figure 1a-c). The negative control region (240,085,545 – 240,101,752; *ρ* = 3 windows) showed no difference in the odds of a non-zero count between the two groups (*p* = 0.02, Figure 1d). Sequencing read counts averaged higher in pregnant samples for all regions of interest, showing greater levels of hypermethylated *RASSF1* (*p* < 0.001), *STAT5A* (*p* < 0.001), and *ERG* (*p* < 0.001) promoter DNA in pregnant samples (Figure 1e-g), consistent with hypermethylation of these regions only in placental tissue. The negative control region showed a slight increase in average read counts in pregnant samples compared to non-pregnant samples (*p* = 0.007, Figure 1h). The elevated levels of DNA fragments hypermethylated in the placental genome compared to other tissues suggest that cfMeDIP-seq enriched for cell-free DNA of placental origin in pregnant samples. Furthermore, we observed correlations between the gestational age at the time of sample collection and the number of sequencing reads mapping to the *RASSF1* (Pearson’s *r* = 0.39, *p* < 0.001), *STAT5A* (*r* = 0.23, *p* < 0.001), and *ERG* (*r* = 0.23, *p* < 0.001) promoter regions (Figure 1i-k). The negative control region showed a weak correlation between gestational age and read counts (*r* = 0.15, *p* = 0.009, Figure 1l). This further supports that cfMeDIP-seq enriched for placental DNA, as it demonstrates increased enrichment of DNA regions with known hypermethylation in the placenta relative to placental growth.

**Figure 1:**
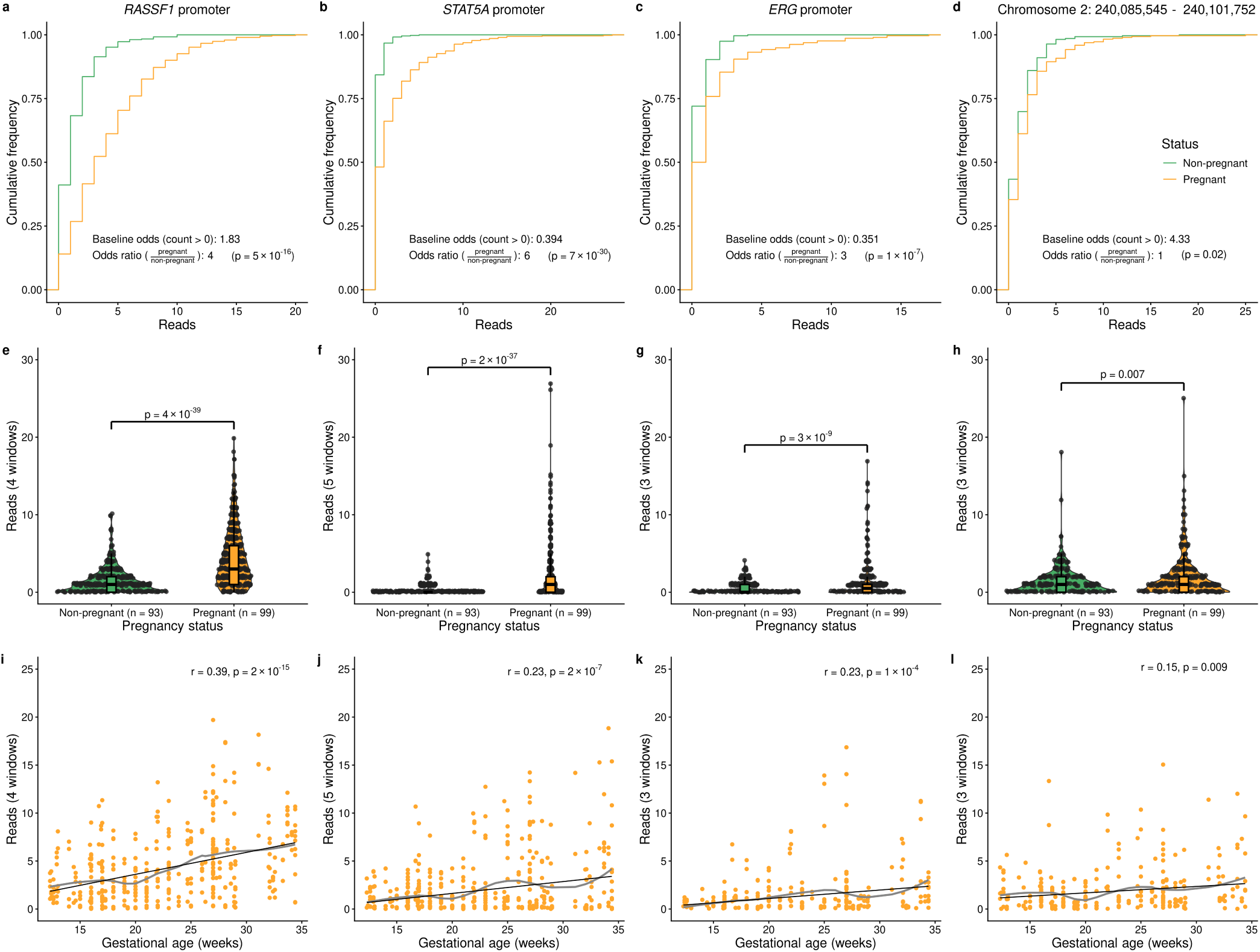
The effect of pregnancy status on the detection by cfMeDIP-seq of for the. **(a)** *RASSF1*, **(b)** *STAT5A*, and **(c)** *ERG* promoters and **(d)** chromosome 2: 240,085,545 – 240,101,752, which exhibits stable methylation across most tissue types. Baseline odds of detection and the increase in odds of detection in the pregnant samples compared to non-pregnant samples appear for each region of interest. For each gene promoter, zero-counts appear most often in non-pregnant samples, while non-zero counts dominate in the pregnant samples. The negative control region shows no difference in the odds of detection between the two groups. **(e)** *RASSF1*, **(f)** *STAT5A*, and **(g)** *ERG* promoter regions exhibit an increase in the average number of read counts in pregnant compared to non-pregnant samples. The negative control region **(h)** showed a similar increase in the pregnant group. Each point represents the read count for an individual 300 bp genomic window, with relative density of reads illustrated by sina and violin plot width. Box plots show median read count and interquartile ranges for each group. P-value determined by a Wilcoxon test. Number of reads mapping to the **(i)** *RASSF1*, **(j)** *STAT5A*, and **(k)** *ERG* promoters and **(l)** negative control region increase across gestational age. Grey line indicates a locally estimated scatterplot smoothing regression of read counts across gestational age, while the black line indicates a linear regression.

Fitting zero inflation hurdle models for each 300 bp window in the datasets revealed that 173,071 out of 536,444 windows (32%) throughout the genome had a significant change in the odds of non-zero read count in pregnant samples (Figure 2). Most of these changes (*ρ* = 166, 785, 96%) reflected a two times or greater increase in the odds of a non-zero read count in pregnant samples.

**Figure 2:**
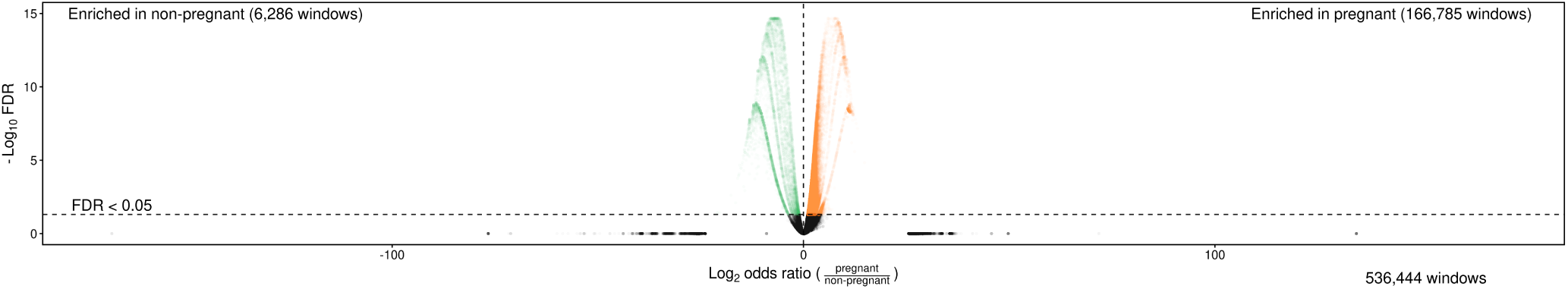
Volcano plot illustrating genomic windows enriched by cfMeDIP-seq in pregnant (*ρ* = 166, 785 windows) and non-pregnant (*ρ* = 6, 286 windows). The x axis denotes the log2 odds ratio of detecting a given read in a pregnant sample relative to a non-pregnant sample. The y axis reflects the –log10 FDR-corrected p-values from the single-window hurdle models.

We observed these increases around the promoter regions of *RASSF1*, *STAT5A*, and *ERG* (Figure 3a-c). The greatest effect usually occurred at or within 600 bp of each gene’s TSS. Nearby regions with low CpG density and known low levels of methylation in placental DNA showed little or no increase in the odds of non-zero read counts. The negative control region showed one 300 bp genomic window with moderate enrichment in pregnant samples (odds ratio = 3) and relative stability across the rest of the region (Figure 3d).

**Figure 3:**
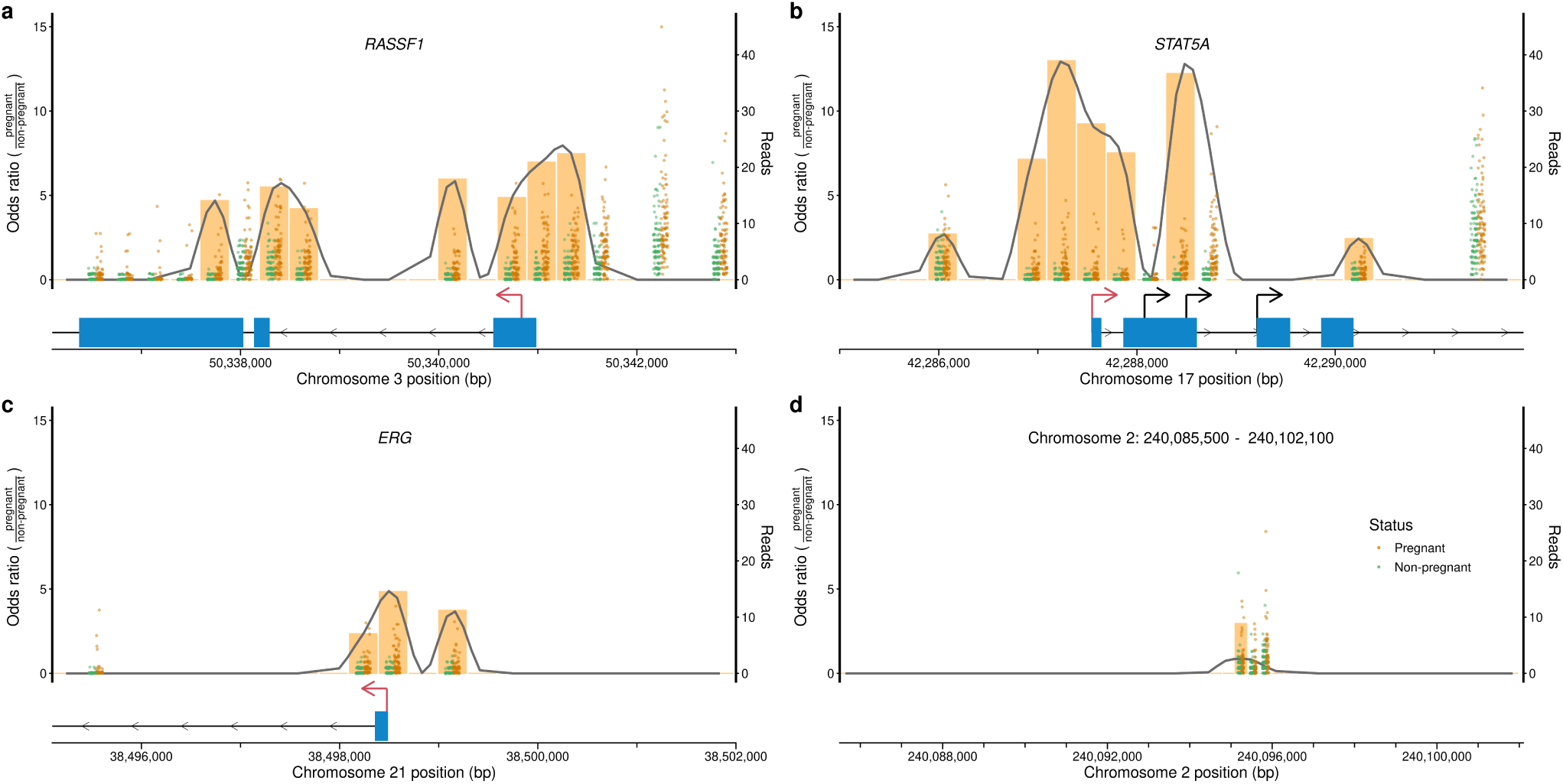
Enrichment of individual 300 bp windows around the TSSs of. **(a)** *RASSF1*, **(b)** *STAT5A*, and **(c)** *ERG*, as indicated by the increase in odds of detection in pregnant samples compared to non-pregnant samples. **(d)** Chromosome 2: 240,085,500 – 240,102,100, which exhibits stable methylation across most tissue types, provides a negative control regions with little expected enrichment. Enrichment peaks at the TSS of each gene with known hypermethylation in placental tissue. Enrichment declines at regions nearby lacking hypermethylation in placental tissue. Individual points denote raw read counts for pregnant and non-pregnant samples at each genomic window. The dashed line indicates a locally estimated scatterplot smoothing regression of enrichment across each genomic region. The blue bar and line segment denote exons and introns of the genes of interest, with TSSs indicated by the arrows above. The red arrow indicates the principal TSS of a given gene.

### IUGR has a distinct cell-free DNA methylation signature

Linear modeling identified 107,505 DMRs in IUGR (*n* = 22) compared to control samples (*n* = 22), adjusting for gestational age, fetal sex, maternal age and BMI, and reads (Equation 1). Most DMRs showed small log2-fold changes, with only 402 DMRs exhibiting a log2-fold change greater than 0.1 in magnitude. No 300 bp genomic windows met statistical significance in pPE (*n* = 14), LOPE (*n* = 23), or PPROM (*n* = 13). Two windows annotating to *CLASP2* and *LINC01992* showed differential DNA methylation between the SPL (*n* = 5) and control samples, while 88,076 windows associated with maternal BMI.

DMRs associated with IUGR mapped to 48,493 repetitive elements, including 51 LINEs, 19,228 Alu elements, 140 non-Alu SINEs, and 3,835 LTRs (Figure 4). Total repetitive elements occurred 0.85 times as frequently as expected based on random sampling of genomic windows (*p* < 0.001). LINEs and non-Alu SINEs showed underrepresentation, occurring 0.61 (*p* < 0.001) and 0.57 (*p* < 0.001) times as often as expected, respectively. LTRs occurred 0.95 times as often as expected (*p* = 0.002). In contrast, Alu elements showed enrichment in the IUGR-associated DMRs, occurring 1.35 times as often as expected from random sampling (*p* < 0.001).

**Figure 4:**
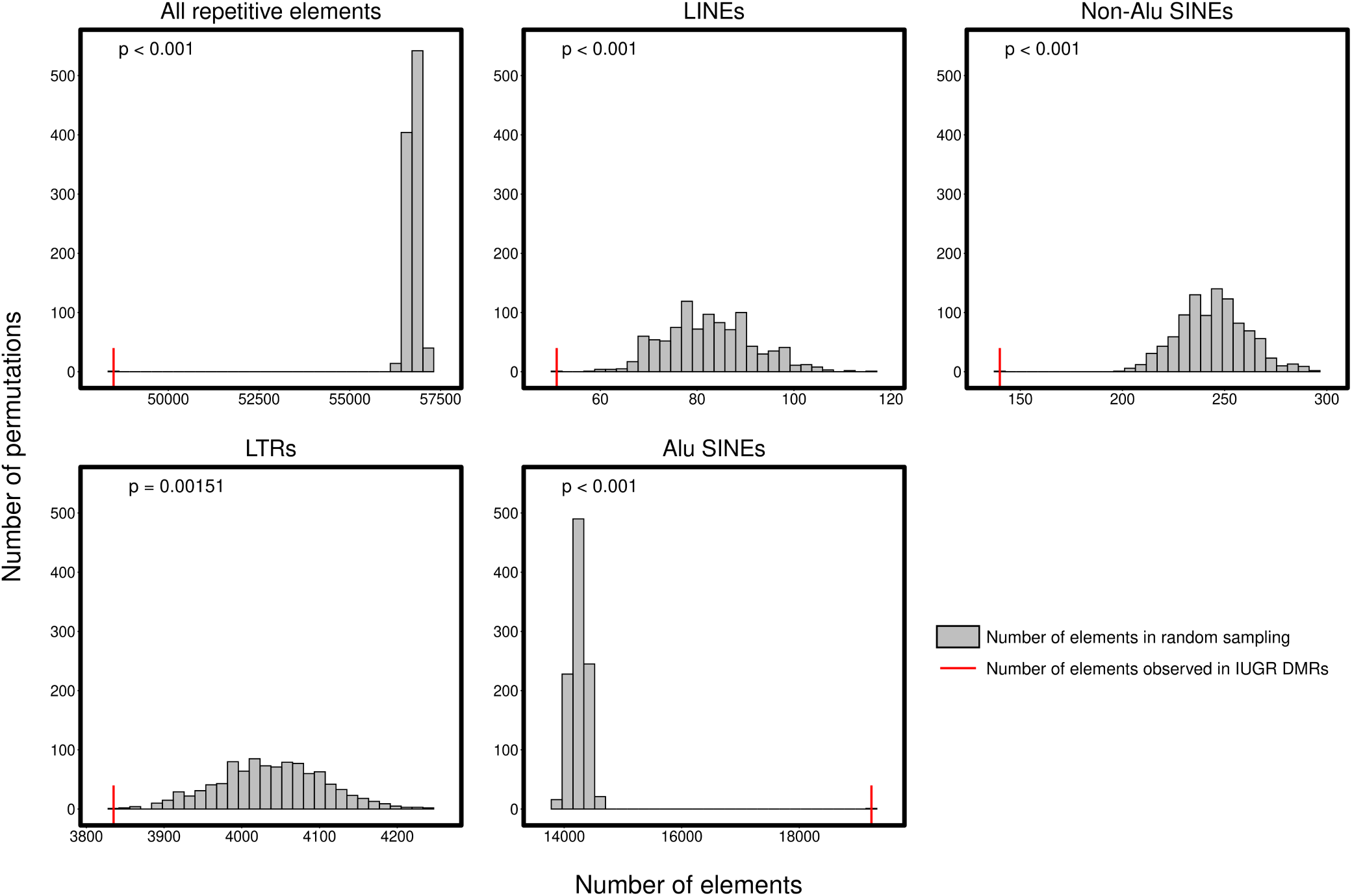
Overall repetitive elements, LINEs, non-Alu SINEs, and LTR retrotransposons exhibited underrepresentation in the set of DMRs relative to a random sampling. Alu SINEs exhibited significant enrichment compared to the expected distribution. The grey histogram illustrates the expected distribution of transposable elements from a random sampling (*X* = 1, 000 permutations) of cfMeDIP-seq windows. The red line denotes the number of each repetitive element observed in the IUGR-associated DMRs.

## Discussion

Placental DNA methylation profiles have shown promise as indicators for placental and overall pregnancy health. The use of cell-free placental DNA bypasses the unnecessary risk of placental tissue sampling during gestation and could serve as the basis of novel, non-invasive risk assessment models for preterm birth and its subtypes. Here, we show that cfMeDIP-seq enriches for cell-free DNA of placental origin. This work opens up new avenues for non-invasive, longitudinal assessment of placental health to predict risk of preterm birth and other pregnancy complications as well as the impact of environmental exposures on pregnancy health. We further report differences in the cell-free DNA methylation signatures of pregnancies resulting in IUGR compared to healthy full-term delivery.

Hypermethylation of the *RASSF1*, *STAT5A*, and *ERG* promoters defines characteristic features of placental DNA (Zhang et al. 2021; Chan et al. 2006; Rahat et al. 2016; Chen et al. 2015). Quantifying hypermethylated *RASSF1* allows calculation of the fraction of cell-free DNA in maternal circulation originating from the placenta (Chan et al. 2006). Our hurdle model analysis showed that regions characteristically hypermethylated in placenta exhibited significantly higher detectability by cfMeDIP-seq in plasma samples from pregnant individuals compared to non-pregnant individuals. Detectability of regions with known stable methylation across tissue types remained unchanged between groups. Nearby genomic regions lacking placental hypermethylation served as further negative control regions where we expected to see no enrichment in pregnant samples. Enrichment near *RASSF1*, *STAT5A*, and *ERG* remained largely confined to the promoter regions, consistent with previously reported hypermethylation in placenta and hypomethylated in other tissues. Nearby regions with no previously reported hypermethylation in placental tissue showed low or lack of enrichment. Non-pregnant samples featured fewer fragments on average from the *RASSF1*, *STAT5A*, and *ERG* promoters, with these regions unlikely to exhibit hypermethylation due to their non-placental origin. As the placenta grows and matures throughout gestation, the amount of placental DNA released into the maternal bloodstream increases (Wang et al. 2013). Our observed correlation between read count at our regions of interest and gestational age aligned with the increased release of cell-free placental DNA associated with placental growth and increased trophoblast turnover. Higher average read counts in pregnant samples and weak correlation with gestational age observed at stably-methylated regions similarly corresponds to increased DNA contribution from the placenta during pregnancy.

Assessment of genome-wide enrichment revealed that roughly one third of genomic regions detected by cfMeDIP-seq showed enrichment in pregnant samples compared to non-pregnant samples. High levels of enrichment tended to correspond to high CpG density, which matches DNA hypermethylation patterns seen in the placental genome (Robinson and Price 2015). The presence of many pregnancy-enriched regions supports that cfMeDIP-seq utility may include assessing placental health and detecting cell-free DNA methylation signatures of complicated pregnancy. Immunoprecipitation of hypermethylated cell-free DNA fragments does not guarantee their placental origin. Fragments of hypermethylated DNA shed from maternal tissues into circulation likely influences DNA methylation signatures. The enriched regions described here offer priority candidate regions for validation of hypermethylation in placental tissue. Methylated DNA immunoprecipitation in combination with techniques such as genetic variation analysis and fragmentomics could accomplish cell-free placental DNA isolation rather than enrichment. However, DNA methylation influence from maternal factors such as hypermethylated hematopoietic cell-free DNA may further strengthen epigenetic signatures of preterm birth by integrating maternal and placental health to indicate pregnancy outcome.

cfMeDIP-seq targets and enriches for hypermethylated placenta-derived cell-free DNA. The inability to examine hypomethylated cell-free placental DNA with cfMeDIP-seq remains a major limitation to characterizing cell-free DNA methylation signatures in their entirety. cfMeDIP-seq does not measure DNA methylation at a single bp resolution and causes loss of DNA methylation information from CpG sparse or partially methylated DNA fragments when normalizing to 300 bp windows. Ongoing work focuses on capturing DNA methylation data for low and partially methylated cell-free placental DNA at higher resolutions to better characterize cell-free placental DNA methylomes throughout pregnancy.

Differential DNA methylation occurs in placental tissue collected at birth from pregnancies complicated by several preterm birth subtypes, with many studies demonstrating differential methylation in preterm preeclamptic placentas (Yuen et al. 2010; Ding 2012; Blair et al. 2013; Anton et al. 2014; Wilson et al. 2018). Our data captured cell-free placental DNA from complicated pregnancies prior to birth, diagnosis, or onset of symptoms. Linear modeling analysis did not reveal any differentially methylated genomic regions in pPE, LOPE, or PPROM. DNA methylation changes associated with these preterm birth subtypes may not yet have occurred at a large enough magnitude to enable detection by cfMeDIP-seq at the gestational ages sampled. Small sample sizes may have resulted in insufficient statistical power to detect small methylation changes for these pathology groups given their heterogeneity. Further, characterizing differential DNA methylation in hypomethylated cell-free placental DNA fragments remains challenging with cfMeDIP-seq, which predominantly captures hypermethylated cell-free DNA fragments. Our findings contribute to the conflicting literature on differential DNA methylation of placenta-derived DNA in preterm and preeclamptic pregnancies, and we further highlight the high degree of heterogeneity within these conditions.

Previous literature has reported differential DNA methylation in placental tissue collected after delivery from IUGR pregnancies (Banister et al. 2011; Ferreira et al. 2011; Koukoura et al. 2012; Wilhelm-Benartzi et al. 2012). When comparing cell-free placental DNA methylation of IUGR samples from prior to symptom onset to control pregnancies, our linear modeling revealed over 100,000 differentially methylated cell-free DNA regions.

We identified an enrichment of Alu elements within IUGR-associated DMRs, whereas LINEs, non-Alu SINEs, and LTRs showed depletion relative to expectation. This patterns likely reflect underlying characteristics of placental DNA methylation, where AluJ and AluS elements retain hypermethylation in the placenta, while other transposable elements typically exhibit stable hypomethylation (Keighley et al. 2023). Reduced overall representation of non-Alu repetitive elements may arise from their hypomethylated state and the enrichment bias of cfMeDIP-seq toward hypermethylation fragments. IUGR-associated methylation changes in these hypomethylated elements may also occur at magnitudes below the detection threshold of cfMeDIP-seq. Prior work has linked altered DNA methylation of transposable elements to growth restriction and preeclampsia, suggesting that dysregulation of normally hypomethylated elements may still contribute to disease despite limited detection in our data (Li et al. 2019; Wilhelm-Benartzi et al. 2012; Keighley et al. 2023). The overrepresentation of Alu elements within DMRs suggests DNA methylation perturbations at these sites that typically undergo methylation-mediated silencing (Keighley et al. 2023).

In conclusion, we have demonstrated that cfMeDIP-seq can enrich for second trimester hypermethylated cell-free placental DNA and characterize cell-free DNA methylation signatures in pregnancies complicated by IUGR. The lack of hypomethylated cell-free placental DNA detection remains a limitation of this technique, however cfMeDIP-seq holds potential in aiding non-invasive assessment of placental health. cfMeDIP-seq utilization could extend to examining how altered cell-free placental DNA methylation relate to functional changes in the placenta and offer novel insights into the impact of pathologies, exposures, and lifestyle factors on placental development and pregnancy health. Our future work will aim to predict the occurrence of preterm birth subtypes based on cell-free DNA methylation signatures throughout gestation, with emphasis on early detection to improve pregnancy outcomes.

## Acknowledgments

This work was supported by the Canadian Health Institutes of Health Research (389866 to M.M.H., postdoctoral fellowship to S.L.W., and a Canada Graduate Research Scholarship—Master’s to K.W.S.), the McLaughlin Centre (MC-2019-07 to M.M.H.), and a Molly Towell Perinatal Research Foundation Fellowship to S.L.W.

## Author Contributions

Conceptualization, S.L.W., M.M.H.; Data curation, K.W.S., N.Y., S.L.W.; Formal analysis, K.W.S., S.L.W.; Funding acquisition, S.L.W., M.M.H.; Methodology, K.W.S., S.G., N.C., P.A., S.V.B., D.D.De C, E.T., S.L.W., M.M.H. Project administration, S.L.W., M.M.H.; Supervision, S.L.W., M.M.H.; Writing — original draft, K.W.S.; Writing — review & editing, K.W.S., N.Y., S.Y.S., S.G., N.C., P.A., T.T., S.V.B., D.D.De C., E.T., S.L.W., M.M.H.

## Data availability

Data for this study will be made available with final publication. All analysis code associated with this project is available at https://github.com/WilsonPregnancyLab/2024cfmedip-iugr.

## Tables and Legends

**Table 1:** Demographics of cohorts.

|  | Non-pregnant | Pregnant |
| --- | --- | --- |
| n | 93 | 99 |
| Age | 65 (5) | 30 (6) |
| BMI | 26.9 (5) | 27.9 (7.1) |
| Total reads | 23,402,398<br>(11,513,869) | 17,357,093<br>(12,316,590) |
| Gestational age (weeks) | — | 23 (6) |
| Fetal sex | — | F: 53 (54%)<br>M: 46 (46%) |

**Table 2:** Demographics of pregnancy cohort.

|  | Control | IUGR | pPE | LOPE | PPROM | SPL |
| --- | --- | --- | --- | --- | --- | --- |
| n | 22 | 22 | 14 | 23 | 13 | 5 |
| Age | 29 (6) | 31 (6) | 31 (5) | 27 (6) | 32 (5) | 29 (6) |
| BMI | 27 (4) | 28 (7) | 27 (5) | 30 (7) | 28 (6) | 32 (18) |
| Total reads | 15,363,990<br>(9,521,795) | 21,607,753<br>(14,472,133) | 21,421,361<br>(14,544,614) | 15,782,163<br>(12,077,563) | 13,155,482<br>(8,370,101) | 14,212,756<br>(13,343,337) |
| Gestational age (weeks) | 23 (7) | 23 (7) | 25 (7) | 20 (6) | 22 (6) | 22 (5) |
| Fetal sex | F: 12 (55%)<br>M: 10 (45%) | F: 13 (59%)<br>M: 9 (41%) | F: 9 (64%)<br>M: 5 (36%) | F: 10 (43%)<br>M: 13 (57%) | F: 7 (54%)<br>M: 6 (46%) | F: 2 (40%)<br>M: 3 (60%) |

## Supplementary Table Titles and Legends

**Table S1. Differentially methylated regions (DMRs) associated with IUGR and their log fold change, p-values, and FDR-corrected p-values.**

## References

1. ’t Oever, R.M. van, Verweij, E.J.T., and Haas, M. de (May 15, 2025). How I use noninvasive prenatal testing for red blood cell and platelet antigens. Blood 145, 2266–2274.

2. Amemiya, H.M., Kundaje, A., and Boyle, A.P. (June 27, 2019). The ENCODE Blacklist: Identification of Problematic Regions of the Genome. Sci Rep 9, 9354. (Visited on 12/14/2024).

3. Anton, L., Brown, A.G., Bartolomei, M.S., and Elovitz, M.A. (June 25, 2014). Differential Methylation of Genes Associated with Cell Adhesion in Preeclamptic Placentas. PLoS ONE 9. C. Oudejans, ed., e100148. (Visited on 12/14/2024).

4. Armengaud, J.B., Yzydorczyk, C., Siddeek, B., Peyter, A.C., and Simeoni, U. (Jan. 2021). Intrauterine growth restriction: Clinical consequences on health and disease at adulthood. Reprod Toxicol 99, 168–176.

5. Ashoor, G., Syngelaki, A., Poon, L.C.Y., Rezende, J.C., and Nicolaides, K.H. (Jan. 2013). Fetal fraction in maternal plasma cell-free DNA at 11-13 weeks’ gestation: relation to maternal and fetal characteristics. Ultrasound Obstet Gynecol 41, 26–32.

6. Banister, C.E., Koestler, D.C., Maccani, M.A., Padbury, J.F., Houseman, E.A., and Marsit, C.J. (July 2011). Infant growth restriction is associated with distinct patterns of DNA methylation in human placentas. Epigenetics 6, 920–927. (Visited on 12/14/2024).

7. Benton, S.J., Leavey, K., Grynspan, D., Cox, B.J., and Bainbridge, S.A. (Dec. 2018). The clinical heterogeneity of preeclampsia is related to both placental gene expression and placental histopathology. American Journal of Obstetrics and Gynecology 219, 604.e1–604.e25. (Visited on 01/12/2025).

8. Bischoff, F.Z., Lewis, D.E., and Simpson, J.L. (2005). Cell-free fetal DNA in maternal blood: kinetics, source and structure. Hum Reprod Update 11, 59–67.

9. Blair, J.D., Yuen, R.K.C., Lim, B.K., McFadden, D.E., Von Dadelszen, P., and Robinson, W.P. (Oct. 1, 2013). Widespread DNA hypomethylation at gene enhancer regions in placentas associated with early-onset pre-eclampsia. Molecular Human Reproduction 19, 697–708. (Visited on 12/14/2024).

10. Bokslag, A., Weissenbruch, M. van, Mol, B.W., and Groot, C.J.M. de (Nov. 2016). Preeclampsia; short and long-term consequences for mother and neonate. Early Hum Dev 102, 47–50.

11. Brockway, H.M., Wilson, S.L., Kallapur, S.G., Buhimschi, C.S., Muglia, L.J., and Jones, H.N. (2023). Characterization of methylation profiles in spontaneous preterm birth placental villous tissue. PLoS One 18, e0279991.

12. Bryzgunova, O.E., Konoshenko, M.Y., and Laktionov, P.P. (Jan. 2021). Concentration of cell-free DNA in different tumor types. Expert Rev Mol Diagn 21, 63–75.

13. Burgener, J.M. et al. (Aug. 1, 2021). Tumor-Naïve Multimodal Profiling of Circulating Tumor DNA in Head and Neck Squamous Cell Carcinoma. Clin Cancer Res 27, 4230–4244.

14. Chan, K.C.A., et al. (Dec. 2006). Hypermethylated RASSF1A in maternal plasma: A universal fetal DNA marker that improves the reliability of noninvasive prenatal diagnosis. Clin Chem 52, 2211–2218.

15. Chen, X., Xiong, L., Zeng, T., Xiao, K., Huang, Y., Guo, H., and Ren, J. (Apr. 15, 2015). Hypermethylated ERG as a cell-free fetal DNA biomarker for non-invasive prenatal testing of Down syndrome. Clin Chim Acta 444, 289–292.

16. Cheng, M.L., Pectasides, E., Hanna, G.J., Parsons, H.A., Choudhury, A.D., and Oxnard, G.R. (Mar. 2021). Circulating tumor DNA in advanced solid tumors: Clinical relevance and future directions. CA Cancer J Clin 71, 176–190.

17. Chu, T., Shaw, P., McClain, L., Simhan, H., and Peters, D. (Jan. 2021). High-resolution epigenomic liquid biopsy for noninvasive phenotyping in pregnancy. Prenat Diagn 41, 61–69.

18. Cindrova-Davies, T. and Sferruzzi-Perri, A.N. (Nov. 2022). Human placental development and function. Semin Cell Dev Biol 131, 66–77.

19. Cragg, J.G. (Sept. 1971). Some Statistical Models for Limited Dependent Variables with Application to the Demand for Durable Goods. Econometrica 39, 829. (Visited on 12/14/2024).

20. Dahlstrøm, B., Romundstad, P., Øian, P., Vatten, L.J., and Eskild, A. (2008). Placenta weight in pre-eclampsia. Acta Obstet Gynecol Scand 87, 608–611.

21. Del Vecchio, G. et al. (June 2021). Cell-free DNA Methylation and Transcriptomic Signature Prediction of Pregnancies with Adverse Outcomes. Epigenetics 16, 642–661.

22. Diaz, L.A. and Bardelli, A. (Feb. 20, 2014). Liquid Biopsies: Genotyping Circulating Tumor DNA. JCO 32, 579–586. (Visited on 12/14/2024).

23. Ding, H.-J. (Apr. 25, 2012). Screening for differential methylation status in human placenta in preeclampsia using a CpG island plus promoter microarray. Int J Mol Med. (Visited on 12/14/2024).

24. Edgar, R., Tan, P.P.C., Portales-Casamar, E., and Pavlidis, P. (2014). Meta-analysis of human methylomes reveals stably methylated sequences surrounding CpG islands associated with high gene expression. Epigenetics Chromatin 7, 28.

25. Fan, H.C., Blumenfeld, Y.J., Chitkara, U., Hudgins, L., and Quake, S.R. (Oct. 21, 2008). Noninvasive diagnosis of fetal aneuploidy by shotgun sequencing DNA from maternal blood. Proc Natl Acad Sci U S A 105, 16266–16271.

26. Ferreira, J.C., Choufani, S., Grafodatskaya, D., Butcher, D.T., Zhao, C., Chitayat, D., Shuman, C., Kingdom, J., Keating, S., and Weksberg, R. (Apr. 2011). *WNT2* promoter methylation in human placenta is associated with low birthweight percentile in the neonate. Epigenetics 6, 440–449. (Visited on 12/14/2024).

27. Goldenberg, R.L., Hauth, J.C., and Andrews, W.W. (May 18, 2000). Intrauterine infection and preterm delivery. N Engl J Med 342, 1500–1507.

28. Goldenberg, R.L., Culhane, J.F., Iams, J.D., and Romero, R. (Jan. 5, 2008). Epidemiology and causes of preterm birth. Lancet 371, 75–84.

29. Grisolia, P. et al. (Oct. 15, 2024). Differential methylation of circulating free DNA assessed through cfMeDiP as a new tool for breast cancer diagnosis and detection of BRCA1/2 mutation. J Transl Med 22, 938.

30. Jensen, T.J., Kim, S.K., Zhu, Z., Chin, C., Gebhard, C., Lu, T., Deciu, C., Van Den Boom, D., and Ehrich, M. (Apr. 15, 2015). Whole genome bisulfite sequencing of cell-free DNA and its cellular contributors uncovers placenta hypomethylated domains. Genome Biol 16, 78. (Visited on 12/14/2024).

31. Karimzadeh, M., Ernst, C., Kundaje, A., and Hoffman, M.M. (Nov. 16, 2018). Umap and Bismap: quantifying genome and methylome mappability. Nucleic Acids Res 46, e120.

32. Keighley, L.M., Lynch-Sutherland, C.F., Almomani, S.N., Eccles, M.R., and Macaulay, E.C. (Sept. 26, 2023). Unveiling the hidden players: The crucial role of transposable elements in the placenta and their potential contribution to pre-eclampsia. Placenta 141, 57–64.

33. Khan, A. et al. (Oct. 4, 2023). The application of epiphenotyping approaches to DNA methylation array studies of the human placenta. Epigenetics & Chromatin 16, 37. (Visited on 12/14/2024).

34. Kirsh, V.A. et al. (Apr. 19, 2023). Cohort Profile: The Ontario Health Study (OHS). International Journal of Epidemiology 52, e137–e151. (Visited on 08/26/2025).

35. Knijnenburg, T.A. et al. (Mar. 19, 2019). Genomic and molecular characterization of preterm birth. Proc Natl Acad Sci U S A 116, 5819–5827.

36. Koukoura, O., Sifakis, S., and Spandidos, D.A. (Apr. 2012). DNA methylation in the human placenta and fetal growth (Review). Molecular Medicine Reports 5, 883–889. (Visited on 12/14/2024).

37. Langmead, B. and Salzberg, S.L. (Apr. 2012). Fast gapped-read alignment with Bowtie 2. Nat Methods 9, 357–359. (Visited on 12/14/2024).

38. Lasseter, K., et al. (Aug. 2020). Plasma cell-free DNA variant analysis compared with methylated DNA analysis in renal cell carcinoma. Genet Med 22, 1366–1373.

39. Leon, L.J., Solanky, N., Stalman, S.E., Demetriou, C., Abu-Amero, S., Stanier, P., Regan, L., and Moore, G.E. (Oct. 2016). A new biological and clinical resource for research into pregnancy complications: The Baby Bio Bank. Placenta 46, 31–37.

40. Li, B. et al. (May 7, 2019). Low Maternal Dietary Folate Alters Retrotranspose by Methylation Regulation in Intrauterine Growth Retardation (IUGR) Fetuses in a Mouse Model. Med Sci Monit 25, 3354–3365.

41. Li, S. et al. (July 11, 2023). Comprehensive tissue deconvolution of cell-free DNA by deep learning for disease diagnosis and monitoring. Proc Natl Acad Sci U S A 120, e2305236120.

42. Lin, S., Leonard, D., Co, M.A., Mukhopadhyay, D., Giri, B., Perger, L., Beeram, M.R., Kuehl, T.J., and Uddin, M.N. (Apr. 2015). Pre-eclampsia has an adverse impact on maternal and fetal health. Translational Research 165, 449–463. (Visited on 12/14/2024).

43. Liu, H.-J., Liu, P.-C., Hua, J., Zhao, Y., and Cao, J. (May 2021). Placental weight and size in relation to fetal growth restriction: a case-control study. J Matern Fetal Neonatal Med 34, 1356–1360.

44. Lu, H. et al. (June 9, 2022). Detection of ovarian cancer using plasma cell-free DNA methylomes. Clin Epigenetics 14, 74

45. Lun, F.M.F., Chiu, R.W.K., Sun, K., Leung, T.Y., Jiang, P., Chan, K.C.A., Sun, H., and Lo, Y.M.D. (Nov. 2013). Noninvasive prenatal methylomic analysis by genomewide bisulfite sequencing of maternal plasma DNA. Clin Chem 59, 1583–1594.

46. Moore, L.D., Le, T., and Fan, G. (Jan. 2013). DNA methylation and its basic function. Neuropsychopharmacology 38, 23–38.

47. Morgan, T.K. (Feb. 2016). Role of the Placenta in Preterm Birth: A Review. Am J Perinatol 33, 258–266.

48. Moutquin, J. (Apr. 2003). Classification and heterogeneity of preterm birth. BJOG: An International Journal of Obstetrics and Gynaecology 110, 30–33. (Visited on 02/12/2025).

49. Mullahy, J. (Dec. 1986). Specification and testing of some modified count data models. Journal of Econometrics 33, 341–365. (Visited on 12/14/2024).

50. Novakovic, B. and Saffery, R. (2013). Placental pseudo-malignancy from a DNA methylation perspective: unanswered questions and future directions. Front Genet 4, 285.

51. Park, B., Khanam, R., Vinayachandran, V., Baqui, A.H., London, S.J., and Biswal, S. (Jan. 1, 2020). Epigenetic biomarkers and preterm birth. Environmental Epigenetics 6. D. Dolinoy, ed., dvaa005. (Visited on 02/13/2025).

52. Quinlan, A.R. and Hall, I.M. (Mar. 15, 2010). BEDTools: a flexible suite of utilities for comparing genomic features. Bioinformatics 26, 841–842.

53. R Core Team (2024). R: A Language and Environment for Statistical Computing (R Foundation for Statistical Computing).

54. Rahat, B., Thakur, S., Bagga, R., and Kaur, J. (Aug. 2016). Epigenetic regulation of STAT5A and its role as fetal DNA epigenetic marker during placental development and dysfunction. Placenta 44, 46–53.

55. Ritchie, M.E., Phipson, B., Wu, D., Hu, Y., Law, C.W., Shi, W., and Smyth, G.K. (Apr. 20, 2015). limma powers differential expression analyses for RNA-sequencing and microarray studies. Nucleic Acids Res 43, e47.

56. Robinson, W.P. and Price, E.M. (Feb. 26, 2015). The human placental methylome. Cold Spring Harb Perspect Med 5, a023044.

57. Salomon, L.J., Sotiriadis, A., Wulff, C.B., Odibo, A., and Akolekar, R. (Oct. 2019). Risk of miscarriage following amniocentesis or chorionic villus sampling: systematic review of literature and updated meta-analysis. Ultrasound in Obstet & Gyne 54, 442–451. (Visited on 07/30/2025).

58. Schneider, V.A. et al. (May 2017). Evaluation of GRCh38 and de novo haploid genome assemblies demonstrates the enduring quality of the reference assembly. Genome Res. 27, 849–864. (Visited on 07/30/2025).

59. Shear, M.A., Swanson, K., Garg, R., Jelin, A.C., Boscardin, J., Norton, M.E., and Sparks, T.N. (Feb. 2023). A systematic review and meta-analysis of cell-free DNA testing for detection of fetal sex chromosome aneuploidy. Prenat Diagn 43, 133–143.

60. Shen, S.Y., Burgener, J.M., Bratman, S.V., and De Carvalho, D.D. (Oct. 2019). Preparation of cfMeDIP-seq libraries for methylome profiling of plasma cell-free DNA. Nat Protoc 14, 2749–2780.

61. Shen, S.Y., et al. (Nov. 2018). Sensitive tumour detection and classification using plasma cell-free DNA methylomes. Nature 563, 579–583. (Visited on 12/14/2024).

62. Steegers, E.A.P., Dadelszen, P. von, Duvekot, J.J., and Pijnenborg, R. (Aug. 21, 2010). Pre-eclampsia. Lancet 376, 631–644.

63. Suhag, A. and Berghella, V. (June 2013). Intrauterine Growth Restriction (IUGR): Etiology and Diagnosis. Curr Obstet Gynecol Rep 2, 102–111. (Visited on 12/14/2024).

64. Sun, K. et al. (Oct. 6, 2015). Plasma DNA tissue mapping by genome-wide methylation sequencing for noninvasive prenatal, cancer, and transplantation assessments. Proc Natl Acad Sci U S A 112, E5503–5512.

65. Wang, E., Batey, A., Struble, C., Musci, T., Song, K., and Oliphant, A. (July 2013). Gestational age and maternal weight effects on fetal cell-free DNA in maternal plasma. Prenatal Diagnosis 33, 662–666. (Visited on 12/15/2024).

66. Wang, X.-M., Tian, F.-Y., Fan, L.-J., Xie, C.-B., Niu, Z.-Z., and Chen, W.-Q. (Jan. 3, 2019). Comparison of DNA methylation profiles associated with spontaneous preterm birth in placenta and cord blood. BMC Med Genomics 12, 1.

67. Wickham, H. (2016). ggplot2 Use R! (Springer International Publishing). (Visited on 12/15/2024).

68. Wilhelm-Benartzi, C.S., Houseman, E.A., Maccani, M.A., Poage, G.M., Koestler, D.C., Langevin, S.M., Gagne, L.A., Banister, C.E., Padbury, J.F., and Marsit, C.J. (Feb. 2012). In Utero Exposures, Infant Growth, and DNA Methylation of Repetitive Elements and Developmentally Related Genes in Human Placenta. Environ Health Perspect 120, 296–302. (Visited on 12/14/2024).

69. Wilson, S.L., Leavey, K., Cox, B.J., and Robinson, W.P. (Jan. 1, 2018). Mining DNA methylation alterations towards a classification of placental pathologies. Human Molecular Genetics 27, 135–146. (Visited on 12/14/2024).

70. Wilson, S.L. and Robinson, W.P. (Apr. 2018). Utility of DNA methylation to assess placental health. Placenta 64, S23–S28. (Visited on 02/12/2025).

71. Wilson, S.L., Shen, S.Y., Harmon, L., Burgener, J.M., Triche, T., Bratman, S.V., De Carvalho, D.D., and Hoffman, M.M. (Sept. 19, 2022). Sensitive and reproducible cell-free methylome quantification with synthetic spike-in controls. Cell Rep Methods 2, 100294.

72. You, Y.-A., Kwon, E.J., Hwang, H.-S., Choi, S.-J., Choi, S.K., and Kim, Y.J. (July 10, 2021). Elevated methylation of the vault RNA2-1 promoter in maternal blood is associated with preterm birth. BMC Genomics 22, 528.

73. Yuen, N., Lemaire, M., and Wilson, S.L. (Dec. 9, 2024). Cell-free placental DNA: What do we really know? PLoS Genet 20. M. Snyder, ed., e1011484. (Visited on 12/14/2024).

74. Yuen, R.K., Peñaherrera, M.S., Von Dadelszen, P., McFadden, D.E., and Robinson, W.P. (Sept. 2010). DNA methylation profiling of human placentas reveals promoter hypomethylation of multiple genes in early-onset preeclampsia. Eur J Hum Genet 18, 1006–1012. (Visited on 12/14/2024).

75. Zhang, L., Wang, C.-M.-Y., Zhou, W.-P., Chen, Q.-P., Zhou, S., Lei, W., Deng, H., Zhang, L., and Liu, G.-C. (Apr. 2021). Dynamic Changes of Fetal-Derived Hypermethylated RASSF1A and Septin 9 Sequences in Maternal Plasma. Reprod Sci 28, 1194–1199.

76. Zuccato, J.A. et al. (Aug. 3, 2023). Cerebrospinal fluid methylome-based liquid biopsies for accurate malignant brain neoplasm classification. Neuro Oncol 25, 1452–1460.

